# Flow cytometry protocols, relative genome size and ploidy levels for 1104 species of non-apomictic angiosperms from the Eastern Alps – a community resource based on the screening of 45,000 samples

**DOI:** 10.64898/2026.01.21.700804

**Authors:** Petr Koutecký, Teresa Zeni, Marianne Magauer, Alžběta Manukjanová, Gabriel Span, Hana Šípková, Jana Vítová, Tomáš Urfus, Filip Kolář, Peter Schönswetter

## Abstract

Flow cytometry provides a reliable and fast method for estimating genome size and ploidy levels in plants. Until recently, most studies employed fresh tissues, which limits the use of the method with samples from remote areas or when an extremely high number of samples needs to be processed in a short time. Although there is growing evidence that silica-dried material can be used for ploidy estimation in some taxa, no flora-wide study has been available so far. Here, we provide methodological aspects of an unprecedented study exploring ploidy variation of non-apomictic angiosperms in the Eastern Alps. We have analysed ca. 45,000 silica-dried samples of 1135 species using flow cytometry with DAPI as stain. We were able to obtain ploidy level information from 1104 (97%) of species. The unsuccessful species included succulent plants of the family *Crassulaceae* (genera *Jovibarba, Rhodiola, Sedum, Sempervivum*), the achlorophyllous parasitic or mycoheterotrophic genera *Orobanche* and *Hypopitis*, and a handful of others. About 80% of samples were successfully analysed using a single ‘universal’ protocol and leaf tissue, while in the remaining species the use of alternative tissues (such as petioles or flowers) and/or protocol modifications were needed (targeting composition of buffers, duration of fixation or staining time or use of alternative buffers). A total of 377 species (34%) included polyploid cytotypes and 179 (16%) species were ploidy-variable. As a community resource, we provide relative genome sizes and ploidy assignments of 1332 cytotypes retrieved from 1104 species along with methodological details (e.g. buffers, standards, analysed plant organs, histogram quality). We believe that this dataset will facilitate future research in particular species as well as in flora-wide investigations of ploidy level variation of the Central European flora in general. We are confident that novel cytotypes of many species will be discovered in other geographic areas, and we would be delighted if the present dataset could serve the botanical community for comparison.

## Introduction

Polyploidy, i.e. the presence of more than two chromosome sets in zygotes and more than one set in gametes, is a common feature in plants. In angiosperms, which is by far the most diverse extant plant group, about 35% of species are polyploid (Wood et al. 2009). Polyploidization is one of the most prominent speciation mechanisms in plants, as it creates an instantaneous reproductive barrier (Husband & Sabara 2004, but see Bartolić et al. 2024). Wood et al. (2009) estimate that ca. 15% of speciation events in angiosperms (and even more in other groups, such as ferns) are directly connected to ploidy level increase. Polyploidization offers evolutionary advantages by enhancing gene redundancy, which buffers against deleterious mutations and provides new genetic material for the evolution of novel functions through subfunctionalization or neofunctionalization (Birchler & Yang 2022). It can also increase heterosis leading to improved vigour and potentially higher fitness (Comai 2005, Otto 2007).

Increased genetic diversity and adaptability in polyploids can facilitate successful colonization of new environments and enable asexual reproduction in the absence of mates (te Beest et al. 2012). Polyploidy can also confer robustness to environmental changes, enhancing survival under stress and offering resistance to pathogens (van den Peer et al. 2021). Genomic studies have shown that most plants underwent at least one, but usually several, rounds of polyploidization in their evolution; polyploidization events can be traced also in the genomes of animals and fungi (Jiao et al. 2011, van de Peer et al. 2017). Despite that, most extant plant species are functionally diploid, and often have low chromosome numbers (an example being 2*n* = 10 in the model species *Arabidopsis thaliana*; Rice et al. 2015). Polyploid genomes undergo progressive reduction of many duplicated regions, a process called diploidization (Wendel et al. 2015, van de Peer et al. 2017). Therefore, in the long-term perspective, polyploidy may be only a temporary phenomenon in the evolutionary history of a plant species. This also implies that the definition of a taxon’s ploidy level is not absolute but rather depends on the phylogenetic context in which it is considered.

The available studies suggest an increasing frequency of polyploidy towards higher latitudes, although the drivers are not clear (Rice et al. 2019). However, in such large-scale studies as well as in the available flora-wide studies (e.g., Šmarda et al. 2019, Chytrý et al. 2021), each species is typically assigned a single ploidy level even though it is widely acknowledged (Rice et al. 2015) that multiple cytotypes are known in about a quarter of plant species. Such intraspecific ploidy variation may then confer differences in genetic variability, genes underlying environmental adaptations, traits or even the ecological niche within a species (Meirmans and Kolář 2025, Vlček et al. 2025, Celestini et al. 2025). Our limited understanding of intraspecific variation in ploidy level is mostly due to the laborious methodology of chromosome counting. Progress in the application of flow cytometry on plant samples over the last two decades (Kron et al. 2007, Loureiro et al. 2010, Galbraith et al. 2021) has enabled us to generate flora-wide datasets with a sufficient number of replicates for each species. Although fresh leaf tissue is the preferred material for flow cytometry, there is growing evidence that silica-dried material can be used in various species for ploidy estimation (e.g. Sonnleitner et al. 2010, Frajman et al. 2015, Kravanja et al. 2025), which overcomes the limitation of the method when a large number of samples from remote places needs to be processed.

Here, our principal aim is to provide a community resource by reporting on methodological aspects of an unprecedented flora-wide study exploring ploidy variation of non-apomictic angiosperms in the Eastern Alps spanning parts of Austria, Italy, Slovenia, Germany and Switzerland. Using flow cytometry of silica-dried material, we have analysed ca. 45,000 samples, collected along 101 elevational transects. While the distribution of polyploids in the Eastern Alps and the underlying factors will be analysed in follow-up studies, the aim of this study is to present (i) the relative genome sizes and ploidy assignments of 1332 cytotypes retrieved from 1104 species. Further, we show that (ii) the vast majority (97%) of species can be analysed from silica-dried samples, and discuss for which taxa this was not possible. Finally, (iii) we provide various methodological details (e.g. buffers, standards, analysed plant organs, histogram quality), with the aim of facilitating future research in particular species as well as in flora-wide investigations of nuclear genome size variation.

## Materials and methods

### Field sampling

We sampled 101 transects positioned on south-exposed mountain slopes in the Eastern Alps during the summers 2021 and 2022. A transect typically consisted of 5 sampling belts, spanning 100 m of elevation each, separated by 150 m. Each complete transect therefore spanned 1100 m of elevation, with the median belt centred at the timberline. In each of the belts, the pool of native (i.e., non-neophytic) angiosperms was sampled as completely as possible from all types of natural or semi-natural vegetation except for aquatic macrophytes. Species from Annex II of the Habitats Directive (in our case *Campanula zoysii* and *Cypripedium calceolus*) and the taxonomically highly intricate, predominantly apomictic genera *Alchemilla, Hieracium, Pilosella* and *Taraxacum* as well as *Rubus* sect. *Rubus* and the *Rosa canina* species group, which could not be exhaustively sampled without the permanent presence of a specialist for each group, were not sampled. For each species, material from different organs of one individual per belt was collected. Samples were put into labelled 8 × 19 cm cellulose bags (tea bags) and dried and stored intermingled with layers of silica gel in air-tight ca. 36 × 56 × 27 cm plastic boxes. Even if much of the silica gel was translocated towards the bottom of the boxes when driving on unpaved roads, this approach proved very efficient for quickly drying large numbers of samples. When the samples were desiccated, silica gel was re-used after drying at ca. 100 °C for several days. As an exception, samples of *Crassulaceae* (*Jovibarba, Sedum, Sempervivum*) were slightly squashed and dried with paper tissue before putting them into tea bags.

The nomenclature in the manuscript and Supplementary Table S1 follows Schratt-Ehrendorfer et al. (2022) and Bartolucci et al. (2018) for the species not included in the former source. However, we also retained selected older synonyms following Fischer et al. (2008) in the Supplementary Table S1 to facilitate the search for taxa.

### Flow cytometry

The relative genome size was measured using flow cytometry. We used Sysmex (formerly Partec) flow cytometers (Sysmex Space, Sysmex/Partec CyFlow Space, Sysmex/Partec CyFlow Space equipped with a CyFlow Space Autoloading Station), using 365 nm LED as light source. The samples were stained with DAPI (4’,6-diamidino-2-phenylindole) due to lower background and lower peak coefficients of variation than the other frequently used dye, propidium iodide (Sliwinska et al. 2022). Since an AT-selective dye was used, results are expressed as relative genome size (RGS), that is the ratio of the mean fluorescence intensity of the sample and the standard G_1_/G_0_ peaks (i.e. the peaks containing nuclei with 2C DNA content).

The sample preparation protocols are described below in detail and indicated in Supplementary Table S1 along with other technical information, such as standards, pooling of samples, and use of the autoloading station. In general, we followed best-practice recommendations outlined in Sliwinska et al. (2022), Temsch et al. (2022), and Koutecký et al. (2023). All samples from the first collection season were analysed individually. After the second season, we used pooled samples of two individuals in selected species under the following conditions: (a) high quality histograms, (b) no missing data (unsuccessful analyses) in the first season, (c) no substantial RGS variation within a ploidy level in the first season, (d) samples from the same transect. If RGS variation in a pooled sample was suspected, the samples were re-analysed individually. We always used internal standardization with one of the following standards (fresh tissue): *Carex acutiformis, Solanum pseudocapsicum, Bellis perennis, Pisum sativum* ‘Ctirad’, *Pisum sativum* ‘Kleine Rheinländerin’, *Chlorophytum comosum, Vicia faba* ‘Inovec’ (see Temsch et al. 2022 for details on the individual standards). We aimed at using one standard for all samples of one genus. However, in large genera with diverse genome sizes (e.g. *Campanula, Cerastium, Festuca, Salix, Saxifraga*), several standards had to be used to avoid peak overlap or more than 4-fold difference between the sample and the standard mean fluorescence. To allow comparison across species, all RGS values were re-calculated to the same standard (i.e. the sample RGS was multiplied by the RGS ratio of the original to the new standard). We measured the RGS ratio (mean of three or more repeats on different days) for all pairs of standards that do not differ more than 4-fold; for more distant pairs, we computed the expected ratio based on standards with intermediate RGS in a cascade-like manner. The re-calculation matrix is presented in Supplementary Table S2. We also re-calculated the RGS of all species to *Bellis perennis* to allow comparison over the entire dataset.

Initially, we tested the ‘universal’ sample preparation protocol with Otto buffers, which are frequently used in plant flow cytometry (i.e., protocol 1C in Doležel et al. 2007, simplified two-step protocol). About 4–25 mm^2^ (depending on the species) of the dried sample and the fresh internal standard was chopped with a sharp razor blade in ice-cold Otto I buffer (0.1 M citric acid monohydrate, 0.5% Tween20), and filtered through a 42 μm nylon mesh (Uhelon 130T, Silk&Progress, Brněnec, Czech Republic). After ca. 5 min the filtrate was stained with Otto II buffer (0.4 M Na_2_HPO_4_.12H_2_O, 0.2% 2-mercaptoethanol, DAPI 4 μg/ml) and analysed after ca. 5 min of staining. If this approach did not yield satisfactory results (i.e., histograms with coefficients of variation (CV) below 5%, low background), different optimizations of the protocol were tested. Frequent modifications included: (a) use of alternative plant tissue, if available (e.g., leaf petioles instead of leaf blades, flowers, seeds; in case of seeds, the embryo peak was scored), (b) addition of 0.1 M HCl to the Otto I buffer (1:1), supplemented with Tween20 to keep its concentration at 0.5% (this corresponds to the ‘lh3’ buffer from Šmarda et al. 2019), (c) addition of 20 μg/ml polyvinylpyrrolidone (PVP), (d) changing incubation time (in Otto I buffer) to short (< 3 min) or prolonged (> 10 min). In some cases, different buffers were tested, especially LB01 buffer (Doležel et al. 2007), with or without PVP.

Depending on the species (amount of material, histogram quality), fluorescence of 3,000– 5,000 particles was recorded. The gain/voltage parameter was adjusted in a way that no peak was located below channel 100 (on the 1,024-channel linear scale), leaving at least 10 ‘empty’ channels before the first peak. During the first measurements of each species, the lower limit of the scale was set such that half of the fluorescence of the putative left-most (lowest-fluorescence) peak was visible to not overlook any peaks; this practice is especially important in plants with endopolyploidy.

A subset of samples, comprising unproblematic species represented by multiple individuals, was analysed using a flow cytometer with autoloading station, which required modified sample preparation. Approximately 3–4 mm^2^ of plant tissue of both the fresh internal standard and the silica-dried sample were inserted into microtubes arranged in a 96-well rack (DNeasy 96 Plant Kit, Qiagen, Hilden, Germany). A tungsten carbide bead with 3 mm diameter (Qiagen) was added to each microtube, followed by the addition of 400 µl ice-cold Otto I buffer. The plant material was ground in microtubes at 30 Hz for ca. 2 min using a TissueLyser II (Qiagen); grinding time was adjusted for each species. Orientation of the rack was reversed after half of the grinding time had elapsed to ensure homogenous grinding. Separation from the debris was achieved by filtering 100 µl of the nuclei suspension through a 42 μm nylon mesh into a 96-well microplate. The suspension was incubated for ca. 15 minutes before staining with 200 µl of Otto II buffer. CyPAD 1.3 software (Sysmex Partec) was used for sample analysis, with the sample volume set to 100 µl. Adjustments of speed and gain/voltage parameters were done similarly as described above, but generally not changed during the run of a plate. Total running time per plate – and consequently the maximum incubation time in Otto II for the final sample – varied between 2 and 3 hours.

Histograms were evaluated using FloMax 2.9 software (Sysmex Partec) or the package flowPloidy (Smith et al. 2018) for R (R Core Team 2024). Mean fluorescence intensities of the G_1_/G_0_ peaks of the sample and the standard – and their CVs – were recorded and the relative genome size (RGS) calculated.

In *Orchidaceae* samples, mean fluorescence intensities of additional peaks were also recorded. Orchids are known for partial endoreduplication. During the endoreduplication cycles, only a part of the genome is replicated; the genome size ratio between the successive peaks in orchid tissues is therefore less than two (Trávníček et al. 2015). Instead of 2C, 4C, 8C, etc., the DNA content of the individual peak is marked as 2C, 2C+P, 2C+3P etc., P being the replicated part of the genome (P < 2C in orchids, while under normal endopolyploidy P = 2C). The ratio between successive peaks is stable and specific for the individual peak pairs (Fig. 1), i.e. it is different between the first (2C) and the second peak (2C+P), the second (2C+P) and the third peak (2C+3P), etc.; the difference is greater, the smaller the value of P is. The height of the peaks (i.e., the number of cells having a particular DNA content) is variable and partly tissue-specific. In leaf tissue, the 2C peak is sometimes missing (Fig. 1), especially in species in which only a small part of the genome is replicated, while it is always present in immature ovaries (Trávníček et al. 2015). Absence of the 2C peak may lead to wrong estimation of the sample RGS. On the other hand, when P is known and at least two peaks are recorded, their specific RGS ratio allows identifying their identity and calculating the RGS of the true 2C peak. We used this approach for analysis of *Orchidaceae* data within our dataset. First, based on samples where ovaries were available (1–52 per species), we recorded the RGS of the individual peaks and estimated the species-specific value of P. We analysed leaf material from the same individuals (up to 20 per species, if available) to verify the calculations. Last, for the samples where only leaves were available, we identified the individual peaks as 2C, 2C+P, 2C+3P etc. based on their RGS ratios, and estimated the 2C RGS, even if the 2C peak was absent. The estimated P values are listed in Supplementary Table S3.

**Fig. 1.**
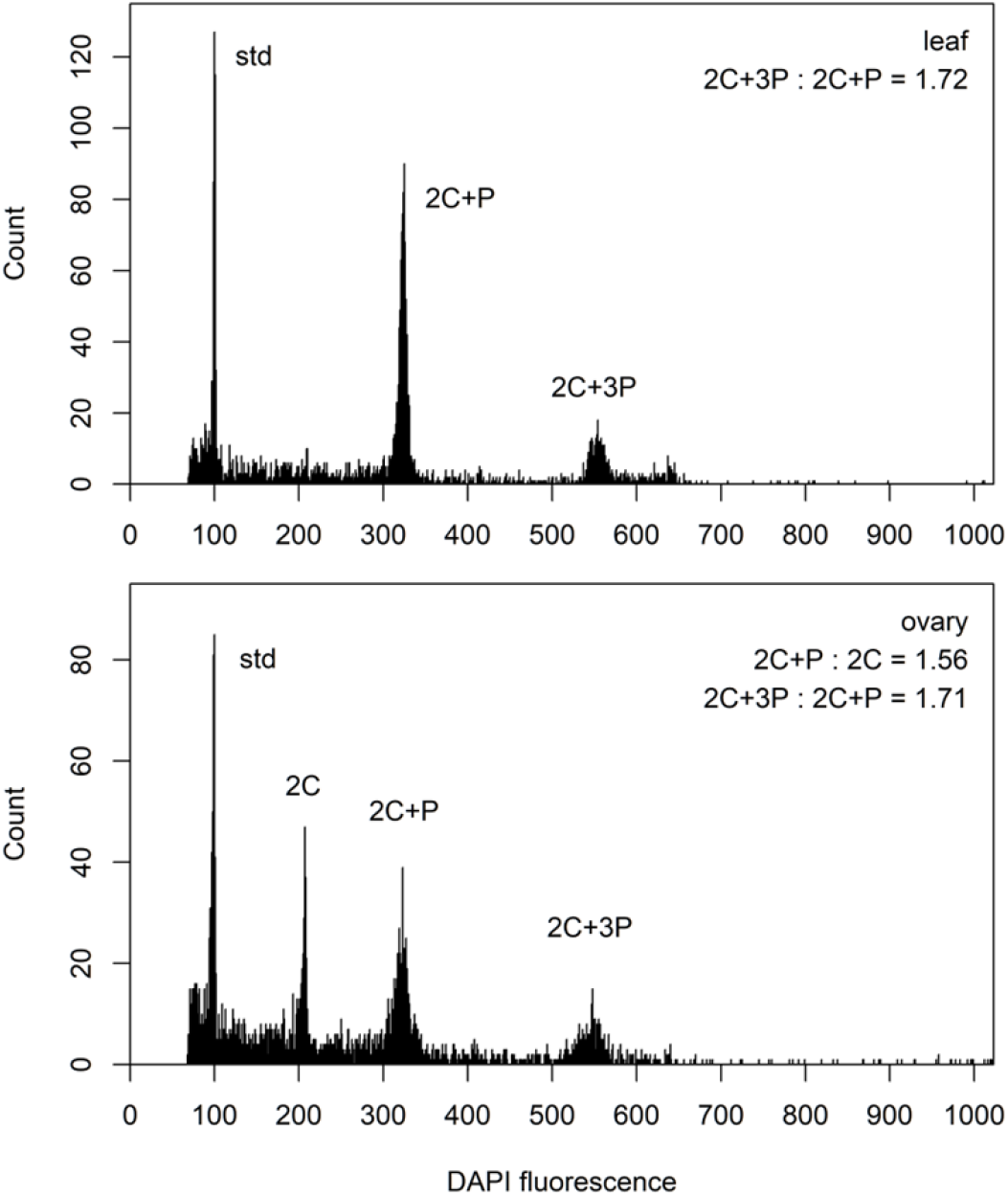
Flow cytometric analysis of a leaf (top) and an ovary (bottom) of the same individual of *Gymnadenia conopsea*. Note the missing 2C peak in the leaf sample and the specific RGS ratios of the successive peaks. *Bellis perennis* was used as the internal standard (std).

Histogram quality was evaluated based on the CV of the sample peak and the amount of background noise; it was ranked in one of three categories (high/medium/low). Additional features were recorded if present, for example the regular presence of multiple peaks due to endopolyploidy or fluorescence shifts due to secondary metabolites (i.e., stable RGS, but both the sample and the standard peaks located at different positions of the fluorescence axis despite the same settings of the cytometer). As a quality check, histograms of the RGS and basic descriptive statistics (mean, standard deviation, coefficient of variation, minimum, maximum) were computed for each species. Outlying RGS values were checked for the analysis quality and possible peak scoring errors; they were re-analysed, if needed. If the outlying RGS was confirmed, we consulted the literature whether karyological/genome size variability is known for a species (e.g. due to aneuploidy or B chromosomes). Further, we checked the plant material for possible determination errors and re-determined the specimen, if necessary. After this procedure, final descriptive statistics was calculated. These statistics are presented in Supplementary Table S1.

### Ploidy assignment

For each species, we extracted the available chromosome counts from the literature and online databases. We preferably used the Austrian chromosome number checklist (Dobeš et Vitek 2000) and online databases and floras of the surrounding countries (Italy: Bedini & Peruzzi 2021–, available at http://bot.biologia.unipi.it/chrobase/index.php; Germany: Paule et al. 2017, available at https://chromosomes.senckenberg.de/item/1; Switzerland: InfoFlora 2025, available at https://www.infoflora.ch; Czechia: Flora of the Czech republic, mainly based on Měsíček & Jarolímová 1992; Slovakia: Marhold et al. 2007, available at https://www.chromosomes.sav.sk/). We also consulted global chromosome databases such as IPCN (IPCN 2025) and CCDB (Rice et al. 2015), the Pladias database of the Czech flora (Chytrý et al. 2021; available at https://pladias.cz/) and specialized literature on individual genera or species; all sources are indicated in Supplementary Table S1.

For each genus, the lowest available chromosome count was considered diploid and we derived the genus’ base chromosome number from it. We preferred records from Europe in general; doubtful unique counts (i.e. those from older literature not documented by photographs or drawings of the chromosome preparations) were disregarded, while reliable extra-European counts were used for alternative ploidy assignment (see below). We assigned ploidy levels to individual species based on their published chromosome counts as well as the measured RGS values. In large and karyologically diverse genera such as *Androsace, Saxifraga* and *Viola*, we applied the same procedure to more homogeneous infra-generic units (subgenera, sections or subsections) instead of the entire genus. In rare cases, our approach may lead to odd-ploidy levels (e.g. 10*x* scored as 5*x*). If no chromosome count was available for a species, we assigned it the same ploidy as the other species of the same genus (subgenus/section) with similar RGS. If the RGS did not match the expected ploidy, we assigned the ploidy based on available taxonomic and karyological literature; all such cases are commented in Supplementary Table S1. In a few cases (24 species, 2%), where two different ploidy assignments (e.g. 2*x* and 4*x* versus 4*x* and 8*x*) are equally likely or when only the extra-European chromosome counts suggested a different base chromosome number, we used the lower ploidy assignments but marked the alternative higher assignments in Supplementary Table S1.

Our approach based on the lowest chromosome count within a genus or a large subgeneric unit yields a more conservative estimate of the frequency of polyploidy compared to the analyses by Rice et al. (2019), who used models of chromosome evolution within an entire genus and scored also diploidized species as polyploids. However, if a species is paleo-polyploid but is (partly) diploidized and thus behaves as a functional diploid (concerning chromosome inheritance, number of copies of most genes, etc.), we preferred to score it as diploid, reserving the polyploid state for evolutionarily young taxa where multiple copies of all chromosomes are still present and likely to segregate.

## Results and discussion

We analysed 1138 species. Three species of *Poa* (*P. alpina, P. pratensis* s. l., *P. nemoralis*) were excluded from the dataset after the first collection season due to the impossibility to link the obtained continuous RGS variation with ploidy levels; most likely, these species are apomictic and frequently aneuploid. Among the remaining 1135 species, RGS and ploidy information was obtained for 1104 species, i.e. 97% (Supplementary Table S1). This is a higher success rate than 87% reported by Suda & Trávníček (2006) based on an analysis of dried tissues of 60 species. The difference may be at least partly caused by testing various buffers and other adjustments of the methodology in our study (see below). Slightly different methods of drying may also contribute to the better performance of the flow cytometry analyses: Suda & Trávníček 2006 used air-drying at 40°C, while we used silica gel at room temperature, which is expected to desiccate the tissues more quickly (Bainard et al. 2011).

The unsuccessful species included succulent plants of the family *Crassulaceae* (all species of the genera *Jovibarba, Rhodiola, Sedum, Sempervivum*), the achlorophyllous parasitic or mycoheterotrophic genera *Orobanche* and *Hypopitis* (syn. *Monotropa*), and a handful of others. We hypothesize that in succulent *Crassulaceae* desiccation was not fast enough to preserve the nuclei. Moreover, plants of this family are fairly recalcitrant even when analysed from fresh material (specific buffers and procedures must be used; Šmarda et al. 2019). The genera *Orobanche* and *Hypopitis* (here, all three analyses of *H. monotropa* were unsuccessful while one out of three analyses of *H. hypophegea* was successful) are somewhat similar to the *Crassulaceae* succulents in having fleshy tissues, in which desiccation might have failed. Among the other unsuccessful species, *Potentilla micrantha* and *Rubus idaeus* belong to the family *Rosaceae* that is known to contain staining inhibitors (Čertner et al. 2022), while we were not able to isolate enough nuclei from most of the *Helianthemum alpestre* samples due to the extremely high amount of mucilage. We have no simple explanation why samples of *Pseudofumaria lutea* did not yield any results. The other species, for which the analyses failed (*Dianthus seguieri, Rumex nivalis, Saxifraga adscendens, Tozzia alpina*) were represented by a single sample each, and the failure might rather be a random event. A low success rate (50% or less) was observed in five additional species, of which only *Galium odoratum* (*Rubiaceae*) and *Pinguicula leptoceras* (*Lentibulariaceae*) were represented by multiple samples, while the other species were represented by two samples each (*Epilobium dodonaei, Saxifraga aspera, Xerolekia speciosissima*)

Almost half of the successfully analysed species (527 out of 1104, 48%) were measured using the standard protocol with Otto buffers, without any specific modifications (such as adjusting buffer composition, incubation time or staining time). Additional 348 species (32%) were measured using a flow cytometer with autoloading station and with Otto buffers used for sample preparation; these samples would also most likely be successfully measured with the standard protocol. This shows that about 80% of the samples were successfully analysed using the standard methodology; this number roughly corresponds to the results of Suda & Trávníček (2006) who obtained acceptable histograms in 52 out of 60 (86.7%) species using the same protocol. In 20% of species in our study, modifications of the protocol were needed. In 7% of species, we adjusted the sample preparation procedure, especially incubation time in Otto I buffer before staining; both very short (< 3 min) and prolonged times (> 10 min) were tested. In the last 13% of species, buffer composition was adjusted (addition of HCl and/or PVP to the Otto I buffer; see Methods for details) or the alternative buffer (LB01) was used, sometimes combined with an adjusted incubation time. All modifications are detailed in Supplementary Table S1. The top five families that required modifications most often (compared to their frequency in the dataset) include *Poaceae, Rosaceae, Salicaceae, Boraginaceae and Onagraceae*.

The analysis quality was scored as high or medium in 1034 species (93.7% of the successful species and 91.1% of all species). In the remaining 70 species, the quality was scored as low or low to medium. Among the species with mostly low quality histograms, there are members of several families listed by Čertner et al. (2022) as problematic for FCM analysis, such as *Boraginaceae* (*Myosotis, Pulmonaria, Sympytum*), *Rosaceae* (*Aruncus dioicus, Crataegus monogyna, Geum urbanum, Rubus saxatilis, Sanguisorba dodecandra*) and *Violaceae* (*Viola*). We observed low quality of histograms also in some *Apiaceae* (*Anthriscus nitidus, Chaerophyllum hirsutum, Ch. villarsii, Coristospermum seguieri*), *Lentibulariaceae* (*Pinguicula*), *Onagraceae* (*Circaea alpina, Epilobium*), *Santalaceae* (*Thesium*) and some other species from different families. The genus *Helianthemum* (*Cistaceae*) appeared extremely challenging due to the high content of mucilage that hampered isolation of a sufficient number of nuclei.

A total of 377 species (34%) included polyploid cytotypes (i.e. they were exclusively polyploid or mixed-ploidy), while 727 (66%) were exclusively diploid. In addition, 37 species were classified as polyploid under the alternative ploidy assignment (see above), raising the estimated frequency of species including polyploids to 36%. These numbers are lower than other estimates. Rice et al. (2019) report 42.9% of polyploids in the ecoregion “Alps conifer and mixed forests” (the spatially relatively limited alpine and subalpine zone being included in this ecoregion, with no separate estimate available), and 43.4% in lower elevations in Central Europe (“Central European mixed forests”). The latter number is nearly identical to 42.7% found in a detailed study of the Czech flora targeting the same ecoregion (Šmarda et al. 2019). The lower estimated polyploid frequency in our sampling probably reflects mainly our more conservative strategy of ploidy assignment.

In total, we recorded 179 (16%) ploidy-variable species. However, based on our sampling, about 70 of them include only rare autopolyploids that occasionally develop in plant populations due to the rare involvement of viable unreduced gametes and are very unlikely to establish as a separate cytotype (e.g. one triploid found among 164 diploids of *Dryas octopetala*). Such individuals are sometimes reported in extensive flow cytometry screenings (e.g. Slovák et al. 2009, Dušková et al. 2010, Kolář et al. 2016). The number of ploidy-variable species is well-comparable to estimates based on global variation of plant ploidy in the CCDB database reporting that 16.2% of plant species harbor intraspecific variation in their ploidy levels (Rice et al. 2015). Given the high sampling effort for many species in our dataset (for 482 and 268 species, we sampled at least 20 and 50 accessions, respectively), the number of ploidy variable species in our dataset can still be considered relatively low. As for many species we sampled only a part of their ranges, some of our ploidy-uniform species are uniform in the study area and elevations covered by our dataset, but include other cytotypes elsewhere (e.g. *Arabidopsis arenosa, Astragalus australis, Potentilla caulescens, Teucrium montanum, Urtica dioica*; see Supplementary Table S1 for references). Finally, within four species (*Juncus jacquinii, Pseudorchis albida, Pulsatilla alpina, Thesium alpinum*), we discovered cytotypes that significantly differ in RGS (by 20–38%) but are probably of the same ploidy level as the difference is far less than full multiple.

The use of dried plant tissue may pose problems due to shifts in fluorescence intensity as compared to fresh material. While Suda & Trávníček (2006), using DAPI dye, report fluorescence intensity decrease of several percent after desiccation, Bainard et al. (2011) surprisingly observed mild fluorescence increase in some species using propidium iodide dye. Suda & Trávníček (2006) also observed a negative effect of the age of the material, but concluded that the lifetime of dried material stored at room temperature is between 1–4 years. We compared our data with the extensive dataset of Šmarda et al. (2019), who analysed fresh material and reported sample/standard ratios (i.e. RGS) for DAPI dye. After re-calculating the RGS values to the same standard (see Supplementary Table S2), we identified 496 cytotypes that are most likely common to both studies (similar RGS and ploidy invariable-species or the same chromosome counts / ploidy levels reported from both study areas, the Eastern Alps and Czechia). There is no systematic shift in the RGS values between these two datasets; the difference between mean RGS values from dried material (our study) and fresh material (Šmarda et al. 2019) is within ± 10% for 479 cytotypes (96.6%), with the mean difference being close to zero, −0.06% (considering values for fresh material as 100%). Differences up to 10% were observed when different laboratories/instruments analysing identical material are compared (Doležel et al. 1998, Sliwinska et al. 2022). Since we analysed most of the samples within one year after the collection (exceptionally up to 3 years, in case samples needed to be re-analysed), we did not observe any pronounced effect of aging on the success rate or quality of the analyses. We conclude that for most of the studied species, the use of dried material is suitable for assigning the samples to different ploidy levels.

Our results demonstrate that flow cytometry can be effectively used for ploidy level screening from dried material across a broad diversity of plant species, often using a single universal protocol. This is relevant as ploidy level is increasingly recognised as an evolutionarily important trait, which is regularly scored in phylogenetic, evolutionary or ecological studies. By making a flora-wide database of RGS values, ploidy levels and methodological details available, we aim to advance the knowledge of the Eastern Alpine flora in particular and of the Central European flora in general. We are confident that novel cytotypes of many species will be discovered in other geographic areas, and we would be delighted if the present dataset could serve the botanical community for comparison.

## Acknowledgements

This work was funded by the Austrian Science Fund (FWF, project P34092 to P.S.).

We thank Gaia Bartolomeo, Paola Comper, Valentina Corieri, Alexander Neugebauer, Elias Nitz, Adam Seyr, Angus Wylie, Remo Zeni, and Valentina Zeni for help with field work. We are most grateful to the following students and colleagues for conducting a large part of the flow cytometric work, partly in the frame of their Bachelor or Master Theses: Gaia Bartolomeo, Anna Borràs, Valentina Corieri, Michela Demetz, Theresa Ecker, Julia Feil, Sonja Gschwendtner, Mia Helling, Katharina Hirschbichler, Fenja Köchl, Laura Morass, Alexander Neugebauer, Petra Niederkofler, Stefanie Noggler, Daniela Pfister, Daniela Pirkebner, Maurice Pitour, Nina Puecher, Susanna Ringler, Solange Wetzels (from the University of Innsbruck), Eliška Fenclová, Adéla Varvažovská (from the Charles University in Prague), Eva Holá, and Alena Lepší (from the University of South Bohemia in České Budějovice). We are grateful to gardeners of the Botanical Garden of the University of Innsbruck for providing high-quality standard plants all year round.

## Supplementary data

Supplementary Table S1. Relative genome sizes, flow cytometry methodology and ploidy assignment.

Supplementary Table S2. Recalculation matrix between genome size standards (DAPI staining)

Supplementary Table S3. Relative genome sizes (RGS) of *Orchidaceae* (DAPI staining)

